# Combined application of biochar with nitrogen fertilizer improves soil quality and reduces soil respiration whilst sustaining wheat grain yield in a semiarid environment

**DOI:** 10.1101/2021.02.22.432242

**Authors:** Stephen Yeboah, Wu Jun, Cai Liqun, Patricia Oteng-Darko, Zhang Renzhi

**Affiliations:** College of Resources and Environmental Sciences, Gansu Agricultural University, Lanzhou, P.R. China; Gansu Provincial Key Lab of Aridland Crop Science, Gansu Agricultural University, Lanzhou, P.R. China; CSIR-Crops Research Institute, Kumasi, Ghana

## Abstract

This study was conducted to investigate the effect of biochar, straw and N fertilizer on soil properties, soil respiration and grain yield of spring wheat (*Triticum aestivum* L.) in semi-arid Western Loess Plateau of northwestern China. The two carbon sources (straw and biochar) were applied alone or combined with nitrogen fertilizer (urea, 46% nitrogen [N]), whilst the soil without carbon is made up of nitrogen fertilizer applied at 0, 50 and 100 kg N/ha. The experiment was arranged in a randomized complete block design with three replicates and was conducted in 2014, 2015 and 2016 cropping season. Results showed that the greatest grain yields were found with 100 kg N ha^−1^ fertilization rate under biochar, straw and soils without carbon, but the greatest effect occurred on the biochar amended soils. Biochar amendment produced the greatest grain yield at 1906 kg ha^−1^, followed by straw treated soils at 1643 kg ha^−1^, and soils without carbon the lowest at 1553 kg ha^−1^. This results is supported by the fact that, biochar amended soils (at 0–10 cm) increased soil organic C by 17.14% and 21.65% compared to straw treated soils and soils without carbon respectively. Seasonal soil respirations were between 19.05% and 23.67% lower in BN_100_ compared with SN_50_ and CN_100_. Soil respiration reduced with increasing N fertilization rates under all treatments, but the greatest effect occurred on biochar plots. Biochar amended soils decreased carbon emission by 26.80% and 9.54% compared to straw treated soils and soils without carbon amendment respectively. Increased grain yield and the decreased carbon emission in BN_100_ translated into greater carbon emission efficiency (2.88 kg kg^−1^) which was significantly different compared with the other treatments. Combined application of biochar with 100 kg N ha^−1^ in rainfed spring wheat was a suitable agricultural practice.

## Introduction

Nitrogen fertilization and crop residue retention are known to influence soil organic C dynamics significantly [1]. Research has shown positive [2], negative [3] and neutral or non-significant effects [4] of N-fertilization on soil organic carbon (SOC) stocks. This apparent controversy may be explained by the fact that increased SOC resulting from increased crop biomass returned to soil may be offset by soil C loss from processes such as erosion, runoff and enhanced oxidation of SOM through tillage [5]. Application of synthetic N-fertilizers may stimulate heterotrophic respiration [6] and may therefore reduce the soil organic carbon (SOC) stock. This process is assisted by ammonium-based fertilizers because of increased SOC desorption following soil application. Desorption of SOC leads to increased microbial decomposition of otherwise stable forms of C, which may also increase the risk of losses through leaching [7]. Soil incorporation of crop residues, particularly with high C:N ratio, contributes to maintain soil organic C levels, enhance biological activity, and increase nutrient availability [8]. Increasing SOM is a slow process, particularly in semiarid environments due to naturally low biomass-C inputs [6].

Recent increases in carbon dioxide emissions levels require that management practices conducive to improved soil C sequestration should be undertaken [9]. Soil carbon sequestration through application of recalcitrant C-rich biochar is mentioned as a suitable means to mitigate climate change, improve soil fertility [4], and crop productivity [5]. Biochar is a recalcitrant source of C, which following soil application contributes to slow down the turnover of native SOC [10]. However, these effects have vary significantly depending upon the type of biochar used and the soil conditions under which the material is applied.

The Loess Plateau is considered the cradle of agricultural cultivation in China and is widely used for grain production. Yet little attention has been given to developing agro-system specific strategies to soil degradation from these regions. This region has experienced a progressive decline in crop productivity because of soil degradation processes. Progressive loss of soil organic matter, associated with traditional methods of soil cultivation often accelerates soil erosion processes, the decline of soil fertility, and loss of soil organic C [11]. Several studies have been conducted to quantify CO_2_ emissions from loess soils in North West (NW) China [12], but there appears to be limited information available about the specific impact of widely-used agronomic practices involving biochar. Quantifying soil respiration and soil quality from rain-fed dryland cropping systems in the Western Loess Plateau is necessary to improve understanding of residue retention practices on environmental quality and crop productivity. In this study, we tested the hypothesis that application of biochar and straw combined with nitrogen fertilizer under rainfed conditions would reduce carbon dioxide emission whiles increasing crop yield.

The objectives of this study were to determine the effect of biochar and straw used alone or combined with fertilizer-N on selected soil chemical properties, grain yield and soil respiration.

## Materials and Methods

### Study site

The study was conducted during the 2014, 2015 and 2016 growing seasons at the Dingxi Experimental Station (35°28′N, 104°44′E, elevation 1971-m above-sea-level) of the Gansu Agricultural University in Northwestern China. The research station is located in the semiarid Western Loess Plateau, which is characterized by step hills and deeply eroded gullies [13]. This area has Aeolian soils, locally known as Huangmian [14], which equate to *Calcaric Cambisols* based on the [15] description. This soil type has a sandy-loam texture and relatively low fertility with pH of ≈ 8.3, soil organic carbon (SOC) ≤8.13 g kg^−1^, and Olsen-P ≤13 mg kg^−1^. This soil type is primarily used for cropping and is the dominant soil in the district. Long-term (annual) rainfall records for Dingxi show an average of 391.9 mm per year; annual evaporation is 1531 mm and aridity is 2.53. Daily maximum temperatures can rise to 38°C in July, while minimum temperatures usually drop to −22°C in January. Annual cumulative temperatures >10°C are 2240°C and annual radiation is 5930 MJ m^−2^, with 2477 h of sunshine. These conditions are representative of those commonly found within agricultural areas of semiarid environments. Prior to the experiment, the site had been occupied by potatoes (*Solanum tuberosum* L.). Seasonal rainfall recorded during the course of the study was 174.6, 252.5 and 239.4 mm in 2014, 2015 and 2016, respectively (S1 Fig).

### Experimental design and treatment description

The experiment involved addition of different carbon (C) sources; namely: biochar and straw, and N fertilizer in the form of urea (46% N) arranged in a randomized block design with 9 treatments and 3 replications. The treatments were: CN_0_ – control (zero-amendment), CN_50_ – 50 kg ha^−1^ N applied each year, CN_100_ – 100 kg ha^−1^ N applied each year, BN_0_ – 15 t ha^−1^ biochar applied in a single dressing in 2014, BN_50_ – 15 t ha^−1^ biochar (single dressing in 2014) + 50 kg ha^−1^ N applied each year, BN_100_ – 15 t ha^−1^ biochar (single dressing in 2014) + 100 kg ha^−1^ N applied each year, SN_0_ – 4.5 t ha^−1^ straw applied in 2014, SN_50_ – 4.5 t ha^−1^ straw in 2014 + 50 kg ha^−1^ N applied each year and SN_100_ – 4.5 t ha^−1^ straw in 2014 + 100 kg ha^−1^ N applied each year. Biochar was evenly spread on the soil surface in March 2014 and incorporated into the soil using a rotary tillage implement to a depth of ≈ 10 cm. The straw used in the experiment was from a previous wheat crop grown at the Experimental Station. In straw–amended plots, the wheat straw from the previous crop was weighted and returned to the original plots immediately after threshing and spread evenly on the soil surface. Biochar and straw were applied at the same quantity based on the straw returned to the soil every year. All the treatments received a blanket application of phosphorus fertilizer which was applied at 45.9 kg ha^−1^ P as ammonium dihydrogen phosphate (12% N, 52% P_2_O_5_). The N and P fertilizers were applied at sowing using a no–tillage seeder and incorporated into the soil to a depth of 20 cm. Spring wheat (*Triticum aestivum* L. *cv*. Dingxi 35) was sown in mid-March at a rate of 188 kg ha^−1^ seeds at 20-cm row spacing, and it was harvested in late July to early August. The plot’s dimensions were 3–m by 6– m and the individual plots were separated by protection rows that were 0.5 m in width.

### Biochar and straw characterization

The biochar used in the experiment is a commercial milled charcoal sourced from a local supplier (Golden Future Agriculture Technology Co., Ltd, Liaoning, China). Biochar was produced from maize straw through pyrolysis at 350–550°C, which can convert 35% of the biomass to biochar in form of granular particles. Based on the protocol described in [16], biochar properties were determined, which included the following: total C using a CN Analyzer (analytikjena; multi N/C, 2100S, Germany), total nitrogen using the Kjeldahl digestion and distillation procedure as described by [17] and pH with pH meter (model: Sartorius PB–10, Germany) using soil to water ratio of 1: 2.5, total ash content, determined after ignition at 720°C in a muffle furnace for 3 h, biochar (bulk) density, determined as mass per unit volume of a biochar sample collected in 260 mm × 220 mm cylindrical containers, and oven-dried to achieve constant weight, and mineral elements composition, which were determined by acid digestion and elemental analysis using atomic adsorption spectroscopy. The Brunauer–Emmett–Teller (BET) method was used to determine biochar surface area. Straw samples were oven-dried at 70°C for 72 h, and milled to pass through a 1-mm sieve. Total C and N contents of the straw, ash content and pH were determined using the same procedure described for biochar. Table 1 shows the chemical characterization of biochar and straw used in the experiment.

**Table 1:** Characterization of biochar and straw used in the study

| Table 1. Characterization of biochar and straw used in the study |  |  |  |  |  |  |  |  |  |  |
| --- | --- | --- | --- | --- | --- | --- | --- | --- | --- | --- |
| Parameter | pH | BD (g cm<sup>-3</sup>) | SA (m<sup>2</sup> g<sup>-1</sup>) | % |  |  |  |  |  |  |
|  |  |  |  | Ca (%) | Mg | K | C | N | P | Ash content |
| Biochar | 9.2 | 0.68 | 8.75 | 0.8 | 0.47 | 0.51 | 53.28 | 1.04 | 0.26 | 25.5 |
| Straw | 6.5 | / | / | 0.53 | 0.04 | 0.47 | 45.05 | 0.94 | 0.08 | 8.9 |

Values are means of n=2.

### Soil sampling, measurements and analyses

Concurrently with soil respiration measurement, soil temperatures were measured at 5, 10, and 15 cm deep using a geo-thermometer inserted into the soil near the chambers. Soil moisture at the 0–5 and 5–10 cm depth intervals was determined by taking a soil core of 5-cm in diameter, and subsequently drying the soil at 105°C for 24 h each time gas fluxes were measured.

Gravimetric water content at the three depth intervals was multiplied by soil bulk density to obtain the volumetric water content, which is expressed in cm^3^ cm^−3^. Soil samples were collected within 3 days prior to planting and immediately after harvest for determination of soil organic carbon. Undisturbed soil samples were collected from three points in each plot using a soil corer (internal diameter: 4.9 cm) for soil C analysis. The soil from the same depth at the 0–10 and 10– 30 cm intervals was bulked and mixed after removal of large plant material. The samples were air-dried, ground to pass 2 mm, and subsequently sub-sampled and ground again to pass 0.25 mm. Soil organic carbon (SOC) in the fine ground samples was determined by the modified Walkley and Black (1934) wet oxidation method [18]. Easily oxidizable carbon (EOC) was determined by oxidation with 333 mmol/L of KMnO_4_ based on [19]. The method of chloroform fumigation and ex-traction (FE) as described by [20] was used to determine the microbial biomass carbon (MBC).

### Soil respiration (Rs)

Soil respiration from soil was measured using an EGM-4 (British PP Systems, Norfolk, UK) portable CO2 analyzer. The emission was measured between 08:00–11:00 h as recommended by [21] so as to capture diurnal patterns of high microbial activity. Three measurements were taken from each plot at each sampling time to reduce the effects of environmental variation and the mean was used for statistical analysis. The Rs (µmol m^2^/s) was converted to Rs (mg m^2^/h) using equation (1), as follows:

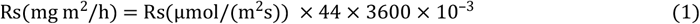

where 44 and 3600 are factors that convert mol to g, and seconds to hour, respectively, and 10^−3^ converts µg to mg.

### Carbon emission (CE)

Carbon emission was estimated based on Rs using the following equation described by [22] and [23]:

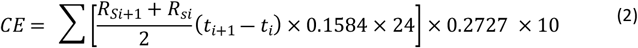

Where: Rs – soil respiration (μmol CO_2_/m^2^/s) measured at biweekly intervals in growing season, i + 1 and i – previous and the current sampling date; t – days after sowing. 0.1584 converted μmol CO_2_/m^2^/s to g CO_2_/m^2^/h, 0.2727 converted g CO_2_/m^2^/h to g C/m^2^/h, and 24 and 10 were to convert g C/m^2^/h to kg/ha for the growing season.

### Carbon emission efficiency (CEE)

To quantify grain yield per unit of carbon emission, carbon emission efficiency (CEE) which was expressed as follows [23]

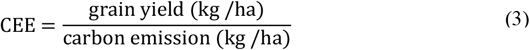

### Grain yield

Plots were hand-harvested using sickles to a height of 5-cm above-the-ground and by discarding the outer edges (0.5-m) from each plot. Grain yield were determined on a dry−weight basis by oven−drying the plant material at 105°C for 45 min and then to constant weight at 85°C [24].

### Statistical analyses

Statistical analyses were undertaken with the Statistical Product Services Solution “22.0’ (IBM Corporation, Chicago, IL, USA) with the treatment as the fixed effect and year as random effect. Differences between-treatments means were determined using Tukey’s honestly significant (HSD) difference test. Significance were determined using a probability level of 5%.

## Results and discussion

### Seasonal variations in soil temperature and moisture

Seasonal temperature increased with time after sowing, and produced a major peak on 21 June, 3 July and 20 July in 2014, 2015 and 2016, respectively (Figs 1a-c). It declined after harvest in August of each year. Generally, higher soil temperatures were recorded during the cropping year 2014 (range: 12.7 to 25.0°C, Fig 1a), followed by 2016 (range: 11.8 to 24.3°C, Fig 1c) and 2015 (range: 10.91 to 23.8°C, Fig 1b). Straw and biochar-treated soils had lower soil temperature while higher soil temperatures were generally observed in soils without addition of C. Generally, the highest seasonal soil moisture values were recorded in straw and biochar treated plots (Figs 2a-c). Lower seasonal soil moisture content was recorded in no carbon treated soils in each of the study year.

**Figs 1a-c.**
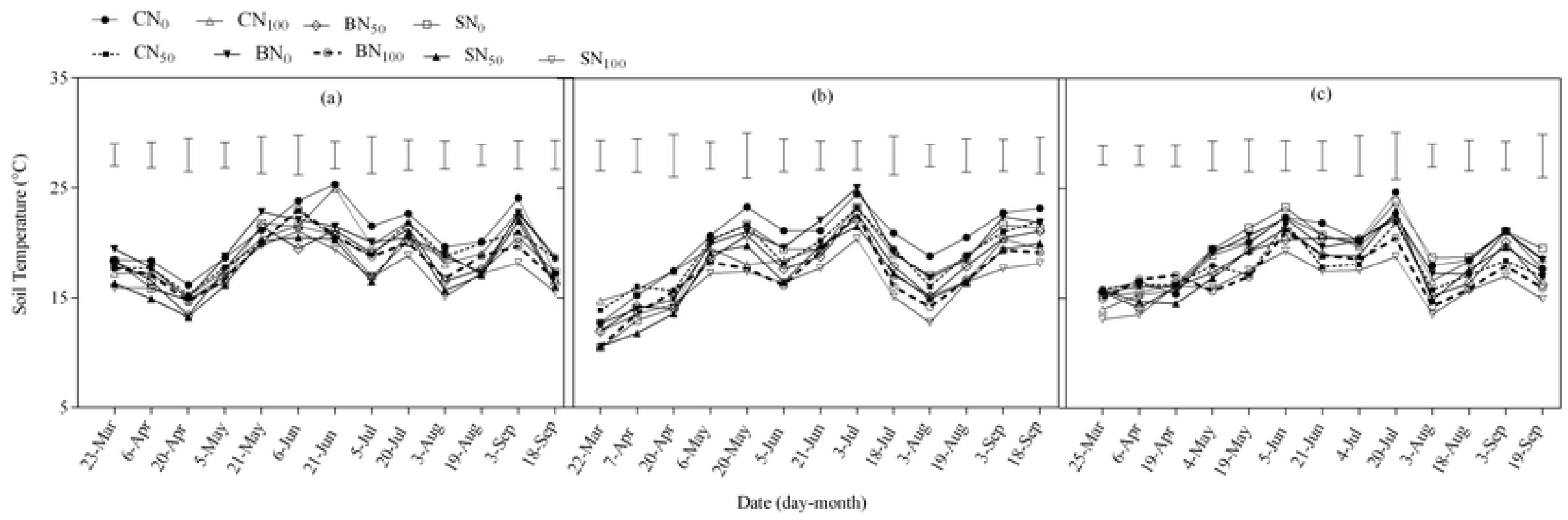
Seasonal soil temperature in 2014 (a), 2015 (b) and 2016 (c) for spring wheat as affected by carbon addition sources. The vertical bars represent the least significant difference (LSD) at p<0.05 among treatments within a measurement date.

**Figs 2a-c.**
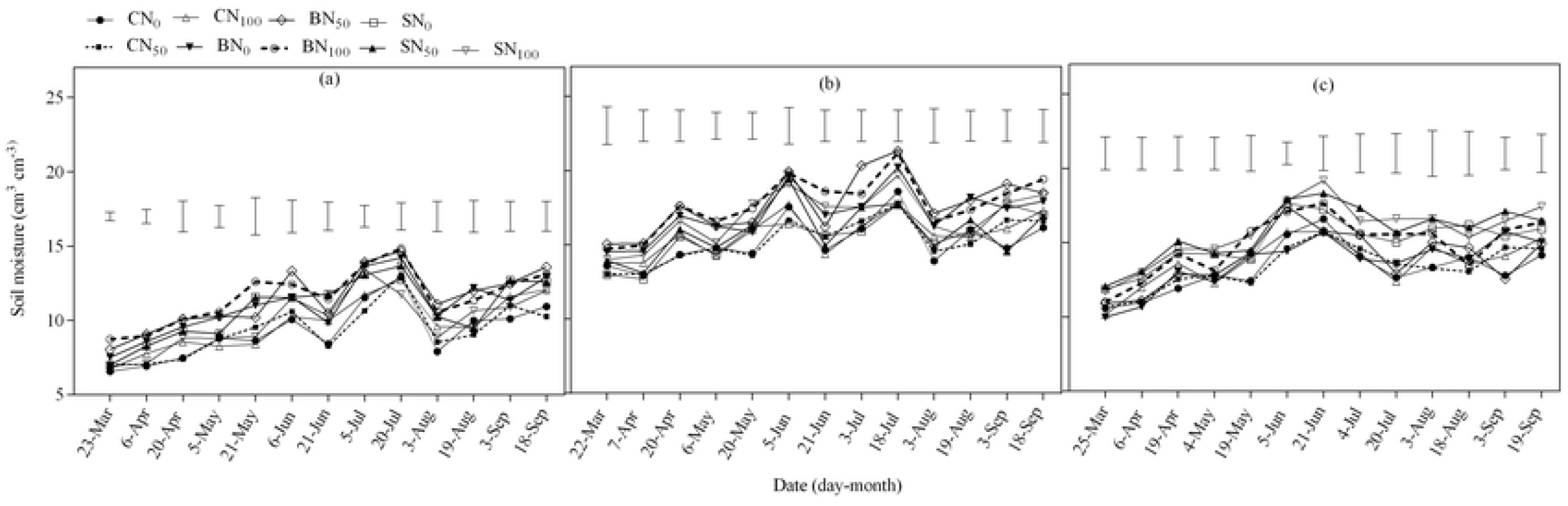
Seasonal soil moisture in 2014 (d), 2015 (e) and 2016 (f) for spring wheat as affected by carbon addition sources. The vertical bars represent the least significant difference (LSD) at p<0.05 among treatments within a measurement date.

According to [11] soil temperature and moisture content, particularly in the 0–30 cm depth interval is important for crop production in dry areas. In the present study, the use of biochar and straw combined with N–fertilizer was shown to increase soil moisture and reduce soil temperature, particularly in the topsoil (0–30 cm depth range). Application of biochar has been reported to increase volumetric water content in soil, improve soil water retention and increase water infiltration [24]. Application of biochar could therefore be used to store more rainfall in soil and increase rainfall use efficiency in dryland areas. Increased soil water holding capacity that follows biochar addition is explained by increased total porosity in soil and specific surface area [25].

### Soil carbon fractions

Carbon, fertilizer-N and year had significant (*P<*0.05) effect on easily oxidizable carbon (EOC), except 10–30 cm where only year had effect (Table 2). There were significant interactions (*P<*0.05) between carbon, nitrogen and year in most of the soil layers reported in here. Both N_50_ and N_100_ fertilization levels significantly affected EOC in different soil layers under all the treatments (Table 3), but the effect was not significant compare to N_0_ in some cases. The N_50_ fertilizer level caused a significant increase of 50.31%, 18.93% and 23.74% on average in EOC at 0–5 cm under biochar soils compare to CN_0_, CN_50_ and CN_100_ respectively. Increasing N fertilization from N_50_ to N_100_ also caused some effect compared to N_0_ but, at a lesser magnitude (Table 3). There was a significant interaction (p<0.05) between C and year for SOC, except at the 10–30 cm depth interval (Table 4). Both carbon and fertilizer-N also had a significant effect on the soil organic C in all the layers studied. The N_50_ and N_100_ fertilizer levels significantly (p< 0.05) enhanced the soil organic carbon under biochar treatments, particularly in 0–5 cm soil layer (Table 4), which corresponded in most cases to significant difference. However, the effect of N_50_ was lesser in many instances relative to N_100_. Irrespective of N level, the soil organic carbon was the lowest with no carbon, followed by straw and then biochar treated soils. Application of N at 50 and 100 kg ha^−1^, increased microbial biomass carbon (MBC) significantly (p< 0.05) in all treatments, but the greatest effect was recorded on straw treated plots, followed closely by the biochar amended soils (Table 5).

**Table 2:**
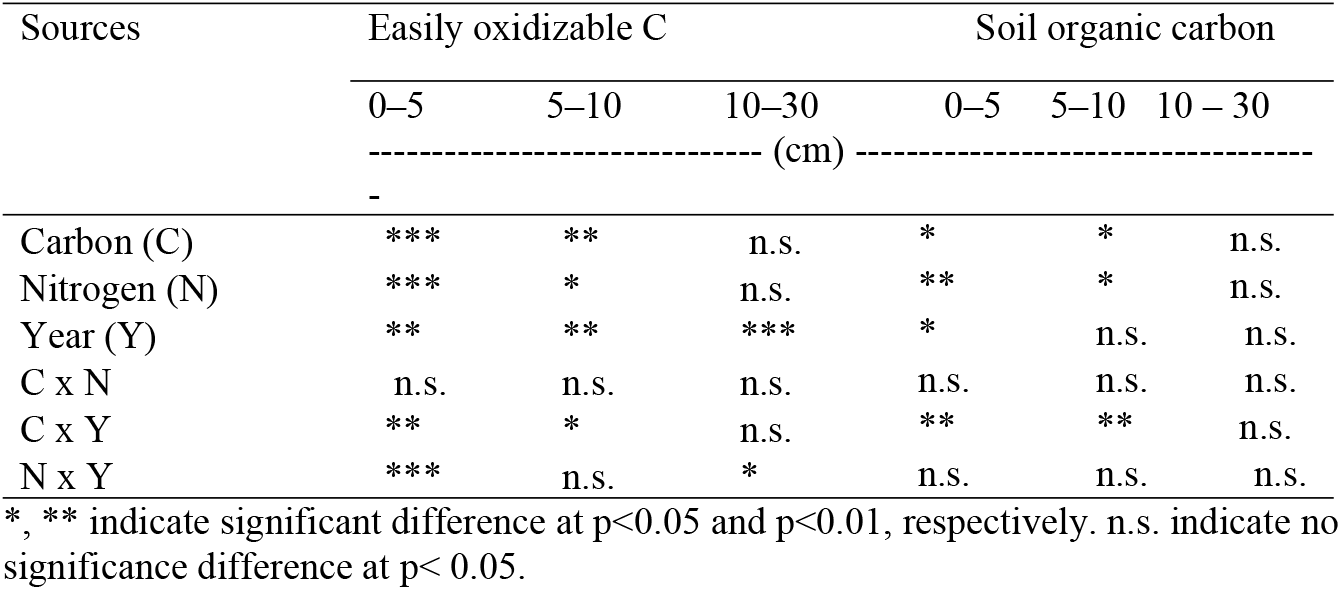
Analysis of variance for carbon, nitrogen and year effects and their interaction

| Sources | Easily oxidizable C |  |  | Soil organic carbon |  |  |
| --- | --- | --- | --- | --- | --- | --- |
|  | 0–5 | 5–10 | 10–30 | 0–5 | 5–10 | 10 – 30 |
|  | ----- (cm) ----- |  |  |  |  |  |
|  | - |  |  |  |  |  |
| Carbon (C) | *** | ** | n.s. | * | * | n.s. |
| Nitrogen (N) | *** | * | n.s. | ** | * | n.s. |
| Year (Y) | ** | ** | *** | * | n.s. | n.s. |
| C x N | n.s. | n.s. | n.s. | n.s. | n.s. | n.s. |
| C x Y | ** | * | n.s. | ** | ** | n.s. |
| N x Y | *** | n.s. | * | n.s. | n.s. | n.s. |

\*, \*\* indicate significant difference at $p < 0.05$ and $p < 0.01$ , respectively. n.s. indicate no significance difference at $p < 0.05$ .

**Table 3:** Easily oxidizable carbon as affected by different treatments

| Treatment |  | Easily oxidizable carbon (g kg<sup>-1</sup>) |  |  |  |  |  |  |  |  |
| --- | --- | --- | --- | --- | --- | --- | --- | --- | --- | --- |
| C source | N rate | 0–5 |  |  | 5–10 |  |  | 10–30 |  |  |
|  |  | ------(cm)----- |  |  |  |  |  |  |  |  |
|  |  | 2014 | 2015 | Mean | 2014 | 2015 | Mean | 2014 | 2015 | Mean |
| No carbon | N<sub>0</sub> | 2.85d | 3.67c | 3.26d | 2.56c | 3.68b | 3.12c | 2.21b | 3.54b | 2.87b |
|  | N<sub>50</sub> | 4.18c | 4.07bc | 4.12c | 3.47bc | 3.96ab | 3.71abc | 3.35ab | 3.57b | 3.46ab |
|  | N<sub>100</sub> | 4.05c | 3.86bc | 3.96c | 3.06bc | 3.86ab | 3.46abc | 2.71ab | 3.65ab | 3.18ab |
| Biochar | N<sub>0</sub> | 4.48bc | 4.16bc | 4.32bc | 3.05bc | 3.77ab | 3.41bc | 2.20b | 3.53b | 2.87b |
|  | N<sub>50</sub> | 5.56a | 4.24bc | 4.90a | 4.95a | 3.91ab | 4.43a | 3.88a | 3.66ab | 3.77ab |
|  | N<sub>100</sub> | 4.68b | 4.24bc | 4.46abc | 3.91abc | 4.23ab | 4.07ab | 2.69ab | 3.90ab | 3.30ab |
| Straw | N<sub>0</sub> | 4.18c | 3.99bc | 4.08c | 3.58abc | 3.70b | 3.64abc | 2.24b | 3.52b | 2.88b |
|  | N<sub>50</sub> | 5.29a | 4.36ab | 4.83ab | 4.13ab | 4.26ab | 4.19a | 2.90ab | 4.14a | 3.52ab |
|  | N<sub>100</sub> | 4.65b | 4.88a | 4.77ab | 3.74abc | 4.47a | 4.10ab | 3.68a | 4.13a | 3.90a |

Values with different letters within a column are significantly different at $p < 0.05$ . $n=3$ .

**Table 4:** Soil organic carbon as affected by different treatments

| Treatment |  | Soil organic C (g kg<sup>-1</sup>) |  |  |  |  |  |  |  |
| --- | --- | --- | --- | --- | --- | --- | --- | --- | --- |
| C source | N rate | 0–10 |  |  |  | 10–30 |  |  |  |
|  |  | 2014 | 2015 | 2016 | Mean | 2014 | 2015 | 2016 | Mean |
| No carbon | N<sub>0</sub> | 9.64c | 9.86c | 10.43e | 9.98d | 9.29b | 9.58b | 9.48e | 9.45c |
|  | N<sub>50</sub> | 10.18bc | 9.92bc | 11.54d | 10.55cd | 10.34ab | 9.73b | 10.71d | 10.26bc |
|  | N<sub>100</sub> | 10.32bc | 10.90bc | 11.70d | 10.97bcd | 9.76ab | 10.10b | 11.05cd | 10.30bc |
| Biochar | N<sub>0</sub> | 11.82ab | 10.28bc | 14.91b | 12.34b | 10.45ab | 10.27b | 12.47b | 11.06b |
|  | N<sub>50</sub> | 12.21ab | 14.04a | 16.01a | 14.09a | 11.59a | 12.66a | 14.41a | 12.89a |
|  | N<sub>100</sub> | 12.42a | 14.09a | 16.26a | 14.26a | 11.04ab | 13.75a | 15.41a | 13.40a |
| Straw | N<sub>0</sub> | 9.71c | 10.14bc | 11.41d | 10.42cd | 9.58ab | 10.12b | 10.69d | 10.13bc |
|  | N<sub>50</sub> | 10.70abc | 10.64bc | 13.77c | 11.70bc | 10.70ab | 10.41b | 11.54bcd | 10.88b |
|  | N<sub>100</sub> | 11.08abc | 11.19b | 14.19bc | 12.15b | 10.92ab | 10.99b | 12.10bc | 11.34b |

Values with different letters within a column are significantly different at $p < 0.05$ . $n=3$ .

**Table 5:** Microbial biomass carbon (MBC) as affected by different treatments

| Treatment |  | MBC (mg kg<sup>-1</sup>) |  |  |
| --- | --- | --- | --- | --- |
| C source | N rate | 0–5 | 5–10 | 10–30 |
| No carbon | N<sub>0</sub> | 164.3f | 157.4h | 118.6f |
|  | N<sub>50</sub> | 190.3d | 177.1fg | 143.0i |
|  | N<sub>100</sub> | 259.4b | 228.3c | 162.8ab |
| Biochar | N<sub>0</sub> | 171.9ef | 167.8gh | 124.2ef |
|  | N<sub>50</sub> | 198.8cd | 184.9ef | 160.2cd |
|  | N<sub>100</sub> | 271.3b | 242.2b | 169.3a |
| Straw | N<sub>0</sub> | 185.3de | 170.2g | 134.9de |
|  | N<sub>50</sub> | 207.9c | 195.3e | 165.4bc |
|  | N<sub>100</sub> | 289.8a | 253.1a | 176.2a |
| **Source** |  |  |  |  |
| Carbon (C) |  | * | * | * |
| Nitrogen (N) |  | * | * | * |
| C X N |  | ns | ns | ns |

267 Values with different letters within a column are significantly different at $p < 0.05$ . $n=3$ .

The significance of retaining crop residues was emphasized in this study by the difference of soil C fractions between the organic amendment soils and the no carbon soils. The greater soil C fractions found in the straw and biochar amended soils supports the hypothesis that increased C inputs results in improved soil carbon stocks. In this study, the increased C content in biochar could be related to its high C content and the fact that biochar could slow down organic C utilization by microbes [26]. The seemingly fast increase in organic C in this study under all treatments could be related to the parent material of the study area. [27] assume soils on calcareous parent material to have a higher ability to store carbon than soils on non–calcareous parent material. The trend observed for microbial biomass carbon (MBC) and easily oxidizable carbon (EOC) was similar to that of organic C. The early response of the carbon fractions to the soil management interventions deployed in this study makes it an important indicator of agronomic sustainability and environmental quality. The mechanism for the increased in MBC in C amended soils could be related to soil residue cover which protects the biological component of the soil and enhance nutrient availability for microbial growth. Such findings are significant to increase crop productivity in nutrient depleted and fragile Loessiah soils.

### Seasonal soil respiration and carbon emission

The seasonal soil respiration over the study period had similar peak times and patterns for all the treatments (Figs 3a-c). The seasonal Rs showed more emission from the straw–mulched treatments in 2014 (Fig 3a) and 2015 (Fig 3b) than from the biochar treated plots and no carbon soils. The major production peaks was observed during the peak growing season of the crop. Significant effects (p<0.05) on seasonal soil respiration were observed on certain occasions of sampling. In most of these occasions, the lowest seasonal soil respiration were observed for BN_100_, and the highest emissions occurred from the CN_100_ and SN_100_ plots. Carbon, N and year showed an interaction effect on carbon emission (Table 6). BN_100_ treatment decreased carbon emissions by 12.37% and 18.80% compared to CN_0_ and CN_50_ soils, respectively (Table 7). Similarly, it decreased carbon emission by 18.18, 33.25 and 32.04% versus SN_0_, SN_50_ and SN_100_ respectively.

**Table 6:** Analysis of variance for carbon, nitrogen and year effects and their interaction

| Source | Rs | CE | CEE | Grain yield |
| --- | --- | --- | --- | --- |
| Carbon (C) | * | *** | *** | ** |
| Nitrogen (N) | ** | *** | *** | ** |
| Year (Y) | * | *** | *** | ** |
| C x N | n.s. | *** | *** | n.s. |
| C x Y | * | *** | *** | n.s. |
| N x Y | n.s. | ** | ** | ** |

\*, \*\* indicate significant difference at $p < 0.05$ and $p < 0.01$ , respectively. n.s. indicate no significance difference at $p < 0.05$ . Rs: Soil respiration; CE: Carbon emission and CEE: Carbon emission efficiency

**Table 7:**
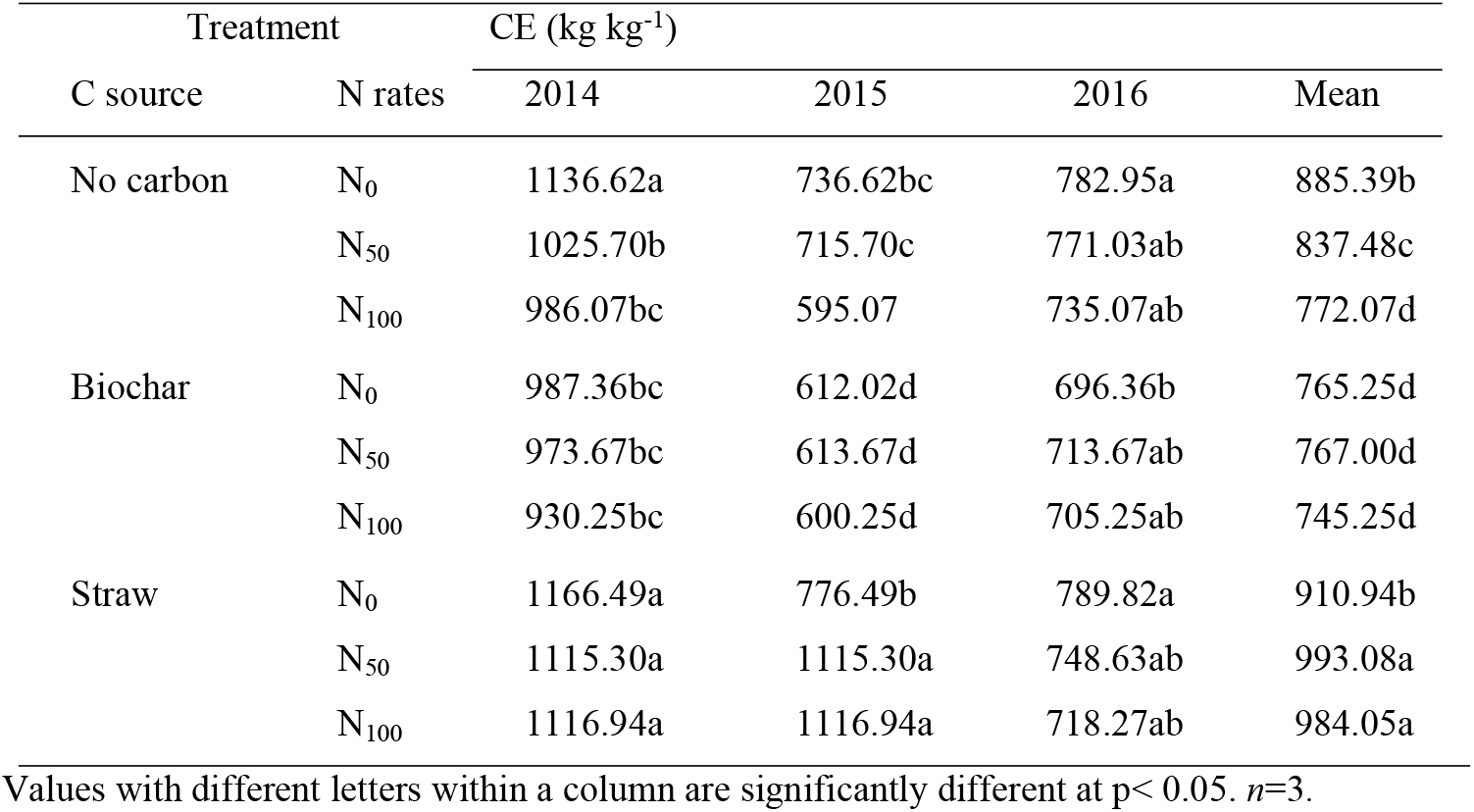
Carbon emission (CE) of spring wheat as affected by different treatments

| Treatment |  | CE (kg kg<sup>-1</sup>) |  |  |  |
| --- | --- | --- | --- | --- | --- |
| C source | N rates | 2014 | 2015 | 2016 | Mean |
| No carbon | N<sub>0</sub> | 1136.62a | 736.62bc | 782.95a | 885.39b |
|  | N<sub>50</sub> | 1025.70b | 715.70c | 771.03ab | 837.48c |
|  | N<sub>100</sub> | 986.07bc | 595.07 | 735.07ab | 772.07d |
| Biochar | N<sub>0</sub> | 987.36bc | 612.02d | 696.36b | 765.25d |
|  | N<sub>50</sub> | 973.67bc | 613.67d | 713.67ab | 767.00d |
|  | N<sub>100</sub> | 930.25bc | 600.25d | 705.25ab | 745.25d |
| Straw | N<sub>0</sub> | 1166.49a | 776.49b | 789.82a | 910.94b |
|  | N<sub>50</sub> | 1115.30a | 1115.30a | 748.63ab | 993.08a |
|  | N<sub>100</sub> | 1116.94a | 1116.94a | 718.27ab | 984.05a |

Values with different letters within a column are significantly different at $p < 0.05$ . $n=3$ .

**Fig 3.**
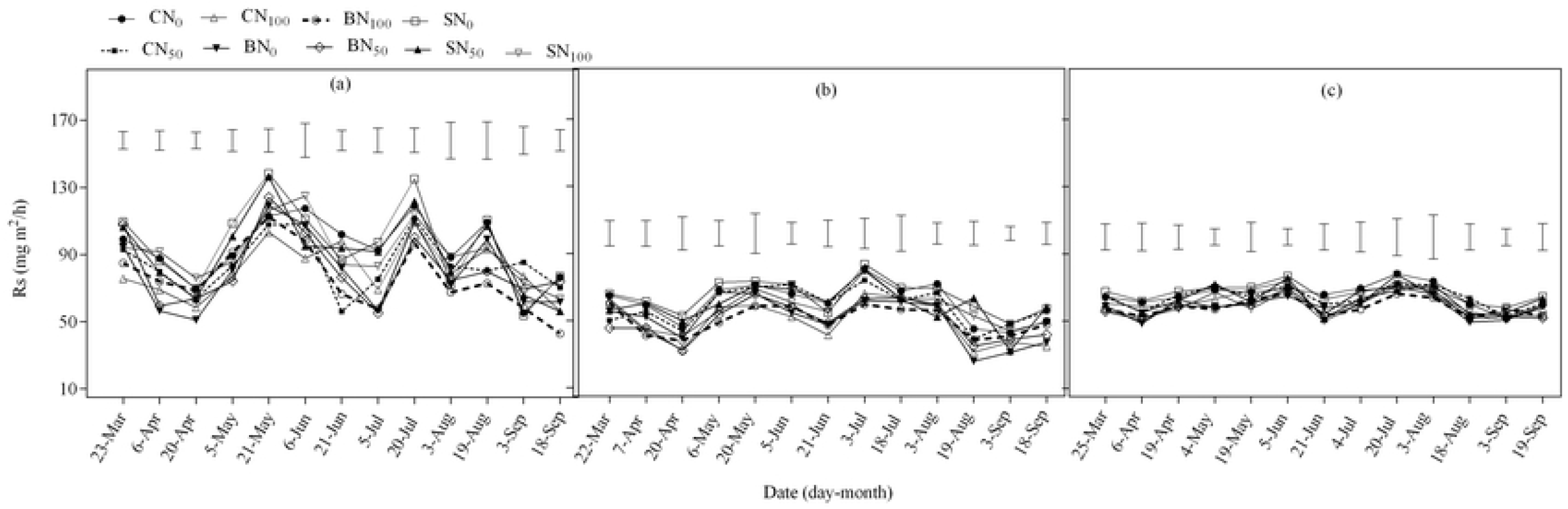
Seasonal soil respiration in spring wheat in 2014 (a), 2015 (b) and 2016 (c) as affected by carbon addition sources. The vertical bars represent the least significant difference (LSD) at p<0.05 among treatments within a measurement date.

All treatments exhibited a similar peak in seasonal soil respiration with a major peak occurring during the peak growing season of the crop, whereas the minor production peak occurred in the least growing period. The peaks period observed in this study was consistent with the pattern of soil temperatures and moisture at the study site [24]. Similar findings regarding variations in Rs with changes in soil temperatures and crop growth have been reported for rice crops [27], spring wheat and field pea crops [28]. Effects of biochar addition on soil respiration have been inconsistent and the underlying mechanisms are still unclear [29]. Both increased and decreased carbon emission has been observed following biochar additions to soils [30]. In this study, the carbon emission was lower in biochar treated soils compared to the other treatments tested. These results were in close agreement with data from [31], who observed that biochar amendments suppressed microbial respiration when applied with N-fertilizer. These responses were attributed to decreased phenol-oxidase activity resulting from N-suppression of white-rot fungi. Moreover, the reductions in carbon when biochar was applied with nitrogen could be attributed to the improved soil organic carbon content. The application of biochar to agriculture acts more as a sink rather than a source because biochar exert high carbon (C) recalcitrant against microbial decay [10]. [26] found reduction in carbon emission following biochar application.

The authors attributed this to the fact that biochar contained high C content and could also protect organic C from utilization. Application of straw caused an overall increase in carbon emission compared to the other treatments over the study period. The higher increase in organic carbon inputs due to the incorporation of wheat straw might explain the larger soil respiration in straw–amended soils in the initial years of the study, which could be associated with the increased availability of C and N substrates, and microbial activities following straw addition [27]. These results suggested that the use of biochar has the potential to improve the overall conditions of semiarid soils, particularly when combined with N fertilizers, without significant increase in soil respiration.

### Grain yield and carbon emission efficiency

Carbon and nitrogen independently affected grain yield of spring wheat (Table 8), and interaction between nitrogen and year were also significant (p<0.05). Application of N_100_ treatment significantly enhanced grain yield of spring wheat under all treatments (Table 8). But the greatest effect was observed on biochar treated soils. The grain yield under N_100_ fertilization significantly increased (p<0.05) by 35.63%, 28.47% and 13.38% under no carbon soils; 33.44%, 37.28% and 39.41% under bio-char soils, and 31.49%, 31.48% and 24.21% under straw soils in 2014, 2015 and 2016 respectively compared to their corresponding N_0_ soils. In this study, nitrogen fertilizer and biochar applied together increased grain yield [9], reflecting the potential of biochar to improve the efficiency with which plants use N-fertilizer. A number of mechanisms have been ascribed to increased crop productivity when biochar is applied in combination with N–fertilizer. [32] noted that improvements to the habitat for beneficial soil microbes are the most likely causes of productivity improvements associated with the application of biochar and inorganic fertilizer. However, other authors [33] have reported that, when biochar and inorganic fertilizers are applied together, an increased nutrient supply to plants may be the most important factor in increasing crop yields. In the current study, increased yield may be attributed to increased nutrient availability as reported in earlier work [34]. Biochar amendments have previously been shown to increase crop productivity by improving soil quality [35]. This study evidenced a positive effect of biochar combined with N fertilizer amendment on soil quality and spring wheat yield consistent over three years.

**Table 8:** Grain yield of spring wheat as affected by different treatments

| Treatment |  |  |  |  |  |
| --- | --- | --- | --- | --- | --- |
| C source | N rates | Grain yield (kg ha<sup>-1</sup>) |  |  |  |
|  |  | 2014 | 2015 | 2016 | Mean |
| No carbon | N<sub>0</sub> | 1305d | 1500d | 1009d | 1271d |
|  | N<sub>50</sub> | 1538cd | 1896bc | 1043cd | 1492bcd |
|  | N<sub>100</sub> | 1770abc | 1927bc | 1144cd | 1614bcd |
| Biochar | N<sub>0</sub> | 1603bcd | 1789cd | 1124cd | 1505bcd |
|  | N<sub>50</sub> | 1905abc | 2133b | 1233bc | 1757abc |
|  | N<sub>100</sub> | 2139a | 2456a | 1567a | 2054a |
| Straw | N<sub>0</sub> | 1502cd | 1658cd | 1111cd | 1424cd |
|  | N<sub>50</sub> | 1852abc | 1944bc | 1182cd | 1659bc |
|  | N<sub>100</sub> | 1975ab | 2180ab | 1380ab | 1845ab |

Values with different letters within a column are significantly different at $p < 0.05$ . $n=3$ .

Carbon emission efficiency (CEE) determines how much grain yield was associated with per unit of carbon emitted. The biochar and straw treated soil improved CEE 32.60 and 7.18% compared the no carbon treated soils (Table 9). The greatest improvement in CEE occurred in all the treatments when N was applied at 100 kg N ha^−1^. The increase in those treatments could be attributed to the higher grain yield and reduction in carbon emission observed in this study. [36] observed increased carbon emission efficiency and attributed it largely to improvement in grain yield. The results suggest biochar + N-fertilizer nutrient management approach can simultaneously manage soil health, improve crop yields while minimizing potential negative impacts of agricultural activities on the environment.

**Table 9:** carbon emission efficiency (CEE) of spring wheat as affected by different treatments

| Treatment |  | CEE (kg kg<sup>-1</sup>) |  |  |  |
| --- | --- | --- | --- | --- | --- |
| C source | N rate | 2014 | 2015 | 2016 | Mean |
| No carbon | N<sub>0</sub> | 1.15e | 2.04e | 1.29e | 1.49g |
|  | N<sub>50</sub> | 1.50d | 2.65d | 1.35de | 1.83e |
|  | N<sub>100</sub> | 1.79bc | 3.24b | 1.56cde | 2.20c |
| Biochar | N<sub>0</sub> | 1.62c | 2.92c | 1.62cd | 2.05d |
|  | N<sub>50</sub> | 1.96b | 3.48b | 1.73bc | 2.39b |
|  | N<sub>100</sub> | 2.30a | 4.09a | 2.24a | 2.88a |
| Straw | N<sub>0</sub> | 1.29de | 2.14e | 1.41de | 1.61f |
|  | N<sub>50</sub> | 1.66c | 1.74f | 1.58cde | 1.66f |
|  | N<sub>100</sub> | 1.77bc | 1.96ef | 1.93b | 1.88e |

Values with different letters within a column are significantly different at $p < 0.05$ . $n=3$ .

## Conclusions

The main conclusions derived from this work are: Application of biochar + N-fertilizer (BN_100_) or straw + N-fertilizer (SN_100_) improved soil organic carbon and microbial biomass carbon, in the 0-5 cm depth interval, to significantly greater extent than the other treatments tested. This translated into higher grain yield in those treatments. Application of biochar + N-fertilizer showed significantly lower carbon emission, but the effect of BN_100_ was consistently greater. Reduced overall emissions and increased yield translated into higher carbon emission efficiency. The results suggest biochar + N-fertilizer nutrient management approach can simultaneously manage soil health, improve yields while minimizing potential negative impacts of agricultural activities on the environment.

## Acknowledgements

Our thanks should go to Gansu Agricultural University for the assistance in the present work. This research was financially supported by the Scientific Research Start-up Funds for Openly-Recruited Doctors (GAU-KYQD-2018-39), the Natural Science Foundation of Gansu province (20JR10RA543), the National Natural Science Foundation of China (41661049,31571594).

## Author Contributions

**Conceptualization:** SY ZR WJ

**Formal analysis:** SY POD

**Funding acquisition:** ZR CL.

**Methodology:** SY ZR CL.

**Software:** POD WJ.

**Writing ± original draft:** SY POD WJ.

**Writing ± review & editing:** SY POD

## Supporting Information

S1 Full precipitation results (XLSX)

S2 Raw Soil data (XLSX)

S3 Fig. Raw soil respiration data (XLSX)

S4 Fig. Raw grain yield data (XLSX)

